# More Insights into the Relation between DNA Ionization Potentials, Single Base Substitutions and Pathogenic Mutations

**DOI:** 10.1101/435354

**Authors:** Fabrizio Pucci, Marianne Rooman

## Abstract

It is nowadays clear that the single base substitutions that occur in the human genome, of which some lead to pathogenic conditions, are non-random and influenced by their flanking nucleobase sequences. However, despite recent progress, the understanding of these “non-local” effects is still far from being achieved. In order to advance this problem, we analyzed the relationship between the base mutability in gene regions and the electron hole transport along the DNA base stacks, as it is one of the mechanisms that have been suggested to contribute to these effects. More precisely, we studied the connection between the observed frequency of single base substitutions and the vertical ionization potential of the base and its flanking sequence, estimated using MP2/6-31G* *ab initio* quantum chemistry calculations. We found a good correlation between the two quantities, whose sign depend on whether SBS is in an exon, an intron or an untranslated region. Interestingly, the correlation appears to be higher for synonymous than for missense mutations, and when considering the flanking sequence of the substituted base in the 3’ rather than in the 5’ direction. A weaker but still statistically significant correlation it found between the ionization potentials and the pathogenicity of the base substitutions. Moreover, pathogenicity is also preferentially associated with larger changes in ionization potentials upon base substitution. With this analysis we gained new insights into the complex biophysical mechanisms that are at the basis of mutagenesis and pathogenicity, and supported the role of electron-hole transport in these matters.

## Background

The understanding of the biophysical mechanisms that drive single DNA base substitutions in different regions of the genome is one of the key questions of the post-genomic era. Indeed, the effects of these mutation processes are potentially responsible for a range of diseases such as cancer and various neurodevelopment disorders. The rationalization of these mechanisms is very complex, since single base substitutions (SBSs) can be triggered by a wide range of factors, *e.g.* chemical species, physical agents or enzymes, through mechanisms such as base deamination, base depurination, or tautomeric shifts. These effects moreover depend on the nucleotide sequence context (1; 2).

The analysis of the large quantity of available genomic data has allowed to firmly state that SBSs do not occur randomly along the genome. For example, one of the well-known mutational signatures observed in cancer genomes occurs at mCpG dinucleotides (3; 4). Methylated cytosines (mC) undergo more frequently spontaneous deamination than unmethylated cytosines, which leads to the C→T transition with higher probability. The second signature, which has been observed in different cancer types, occurs at TpC dinucleotides in which the cytosine is mutated through transition or transversion. It has been related to the overactivity of the APOBEC cytidine deaminases (5; 6). Other signatures are specific to certain types of cancers or diseases (7; 8; 9). For example, the C:G→A:T transversion is a well-known mutation in lung cancer induced by tobacco carcinogens. The “UV signature”, namely C→T mutations at dipyrimidine sites, is caused by ultraviolet light via cyclobutane pyrimidine dimer formation and is commonly found in melanomas.

It became recently clear that there is an important effect of the flanking base sequence on SBSs, which appears to extend beyond the dinucleotide units (2; 10). However, the mechanisms triggering these “non-local” effects are far from clear. Their comprehension is of primary importance and would lead to deep insights into mutagenesis phenomena and the deleteriousness of genetic variations.

In this context, the physical phenomena of electron-hole transfer along the DNA stack comes into play. Exposure to high-energy radiations or to reactive oxygen species generated as by-products of the cellular metabolism can lead to DNA ionization through the creation of electron holes. These holes then migrate along the DNA stack until they remain (more or less) localized in a potential well (12; 14; 15).

Although some electron holes have solely a damaging effect and need to be repaired by specific enzymes (13), others seem to have a positive role as suggested more than a decade ago by the finding that the amount of reactive oxygen species likely to create such holes is regulated by specific proteins and is, for example, higher during cell differentiation (18). The important role played by electron-hole transfer in various biological processes appears increasingly clear (16; 17) and more specifically, in the sequence dependence of the SBSs (2; 14; 15; 25) and their possible pathogenicity (19; 14).

Important steps towards the understanding of long-distance hole transfer in DNA have been achieved. However, large parts of the picture are still lacking. Indeed, experimental measures are challenging due to the complexity of working at the nanoscale (21; 22), and accurate quantum chemistry calculations are extremely heavy for these systems (11; 23).

In this paper, we further analyzed the relationship between the electron-hole transfer along the DNA stack, the sequence-dependence of the mutability and the pathogenicity of the mutations. For that purpose, we calculated the vertical ionization potential (vIP) of all short nucleobase sequences and correlate these to the presence of benign and/or deleterious mutations occurring in cancer and inherited disorders, with the goal of confirming the prominent contribution of hole migration in these matters.

## Results

### Composition of the SBS dataset

We have set up a dataset of single base substitutions which occur in genes, *i.e.* in exons, introns or untranslated regions (UTR), and are annotated as pathogenic, benign, of unclear significance, etc, as described in Methods. In exons, we also distinguish between synonymous and missense variants. Let us start by analyzing which of the four single bases, and which of the nucleobase doublets and triplets mutate significantly more frequently, or less frequently, than expected. The results are summarized in Table 1, and reported exhaustively in Table S3 in the Supplementary Materials.

**Table 1.**
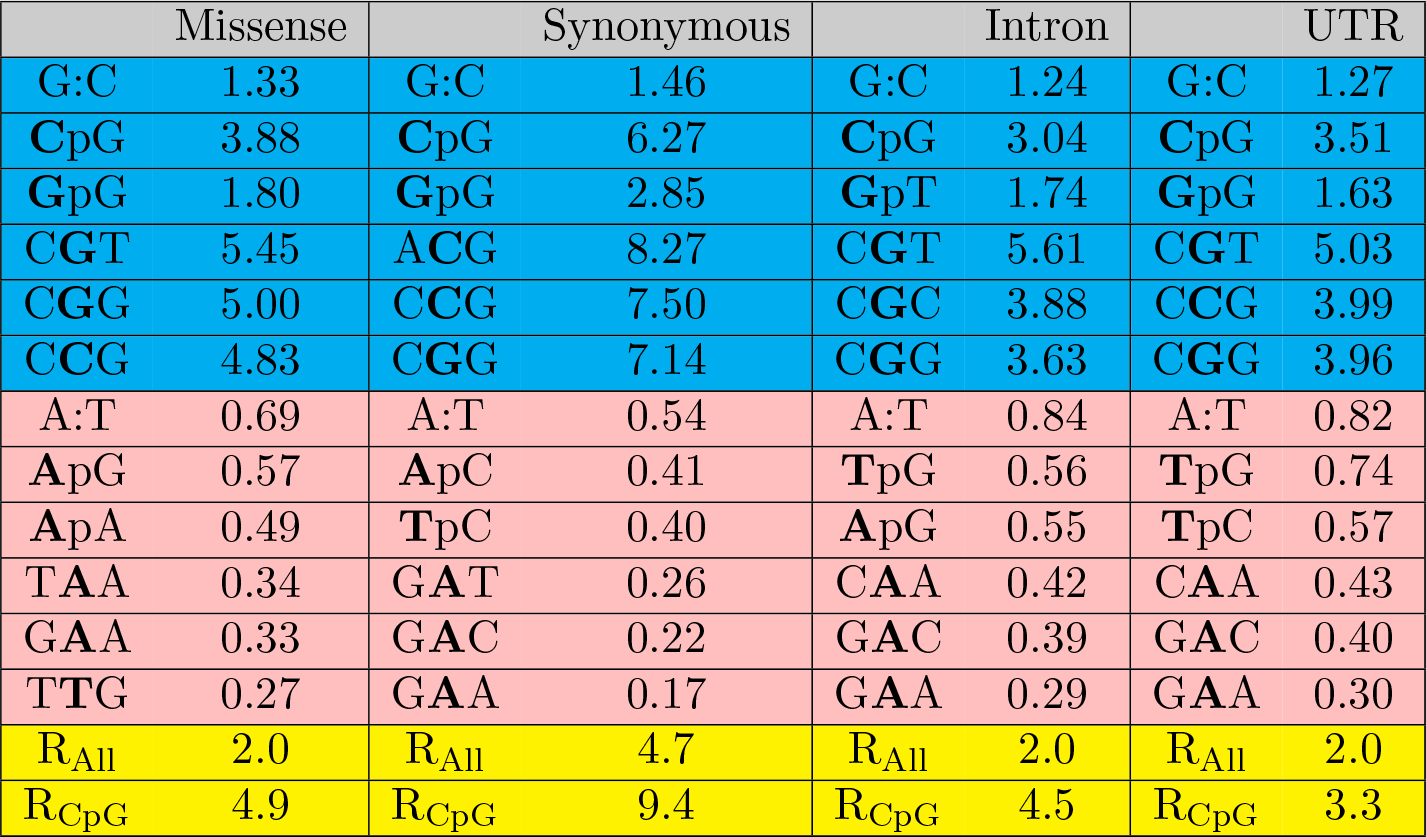
Ratio of the frequencies of mutations in nucleobase motifs computed from the SBS dataset to the frequencies of same motifs obtained from exome data. The nucleotide at which the substitution occurs is in bold. The mutations that are significantly more frequent than expected are on a blue background, and those that are significantly less frequent are on a pink background. R is the ratio of transitions to transversions.

| Missense |  | Synonymous |  | Intron |  | UTR |  |
| --- | --- | --- | --- | --- | --- | --- | --- |
| G:C | 1.33 | G:C | 1.46 | G:C | 1.24 | G:C | 1.27 |
| CpG | 3.88 | CpG | 6.27 | CpG | 3.04 | CpG | 3.51 |
| GpG | 1.80 | GpG | 2.85 | GpT | 1.74 | GpG | 1.63 |
| CGT | 5.45 | ACG | 8.27 | CGT | 5.61 | CGT | 5.03 |
| CGG | 5.00 | CCG | 7.50 | CGC | 3.88 | CCG | 3.99 |
| CCG | 4.83 | CGG | 7.14 | CGG | 3.63 | CGG | 3.96 |
| A:T | 0.69 | A:T | 0.54 | A:T | 0.84 | A:T | 0.82 |
| ApG | 0.57 | ApC | 0.41 | TpG | 0.56 | TpG | 0.74 |
| ApA | 0.49 | TpC | 0.40 | ApG | 0.55 | TpC | 0.57 |
| TAA | 0.34 | GAT | 0.26 | CAA | 0.42 | CAA | 0.43 |
| GAA | 0.33 | GAC | 0.22 | GAC | 0.39 | GAC | 0.40 |
| TTG | 0.27 | GAA | 0.17 | GAA | 0.29 | GAA | 0.30 |
| R<sub>All</sub> | 2.0 | R<sub>All</sub> | 4.7 | R<sub>All</sub> | 2.0 | R<sub>All</sub> | 2.0 |
| R<sub>CpG</sub> | 4.9 | R<sub>CpG</sub> | 9.4 | R<sub>CpG</sub> | 4.5 | R<sub>CpG</sub> | 3.3 |

We clearly observe the well-known bias towards the mutations that occur at C:G base pairs rather than at A:T (25). Indeed, the frequency of mutations of C or G is 1.2 to 1.5 higher than expected on the basis of the frequency of C:G pairs in the human exome. Note that C:G pairs are slightly underrepresented in the human genome (around 40%) but not in the human exome (around 50-55%).

The preference of SBSs to occur on the C base of CpG dinucleotides is even more marked, with a rate that is 3 to 6 times higher than expected from the CpG frequency in the exome (see table S3). Notice also that CpGs are underrepresented in the genome, which can be attributed to the fact that they are the preferential target of DNA-methyltransferase which methylates the cytosine and can then more easily undergo the base substitution mCpG→TpG (25; 26).

The other doublets that are preferentially targeted by mutations are **G**pG in exons and UTRs, and **G**pT in introns, where the substituted base is in bold. In contrast, the least mutated dinucleotides are **T**p(C or G) and **A**p(A, C or G) with a rate that is 6 to 15 times lower than the **C**pG mutation rate. Note that TpA dinucleotides are also underrepresented in all gene regions (26).

Furthermore, SBSs occur preferentially at the middle base of C**G**G and C**C**G triplets in all regions (C**C**G is actually just below the threshold in introns, see Table S3). At the other extreme, G**A**A and some other triplets with A-base substitutions occur at rates that are up to 40 times lower than the C**G**G and C**C**G triplet rates.

An interesting observation is that all trends are systematically amplified for synonymous mutations: the most frequently mutated sequence motifs are always more frequent, and the least frequently mutated motifs are always less frequent, than in missense exonic mutations, in introns and in UTRs. This amplification is moreover quite strong: for **C**pG doublets, for example, the frequency of synonymous mutations is 6.3 times higher than expected, while this value it is only 3.9 for missense mutations. In contrast, the frequencies of missense, intronic and UTR SBSs are quite similar. The reason of the synonymous exception is currently unclear as discussed in the Discussion section.

Another interesting observation is that the preferentially mutated doublets and triplets often differ according to whether they are located in exons, introns or UTRs. This seems to suggest the existence of different mutational mechanisms in these different regions. We will come back to this point in the next sections.

We also observed that transitions constitute 2/3 of the SBSs in the missense, intron and UTR subgroups, and that this frequency increases up to about 5/6 for the synonymous mutations. These values are close to the ones already observed in other analyses (27; 28) where a transition/transversion ratio of 2.3-2.4 was estimated. These ratios substantially increase at **C**pG sites where they reach values of 3-4 for missense, intronic and UTR SBSs, and more than 9 for synonymous mutations.

### Relation between the vIP of base sequence motifs and the frequency of SBSs

An important biophysical mechanism that can be related to the sequence dependence of the SBSs is the presence and migration of radical cations (electron holes) along the DNA molecules. Indeed, due to oxidative stress caused by physical or chemical agents, an electron can be extracted from the DNA. The electron hole then starts migrating along the aromatic rings of the stacked nucleobases, until it remains trapped in a minimum of the ionization potential. The hole can be repaired by specific DNA repair proteins or, if not, trigger a base substitution.

In order to quantify these long-range effects and their relation with SBSs, we started by calculating the vIP of all 4 nucleobases, 16 nucleobase doublets, 64 triplets and 256 quadruplets using M0ller-Plesset perturbation theory as described in Methods. These vIP values, which are directly related to the probability of extraction of an electron from the DNA base-stacking structure, are given in Table S3. We also computed the vIP of all nucleobase quintuplets by taking the mean of the vIPs of the two overlapping quadruplets, and of the sextuplets as the mean of the three overlapping quadruplets. Note that we used the experimental values given in Table S3 for the single-base vIPs, which are slightly different from the calculated ones and yield better results in the present context, as expected.

We then computed the correlations between the vIP of the wild type nucleotides with their flanking base sequences and the frequency of observation of the corresponding sequence segment in the SBS dataset. More precisely, these correlations were calculated for different flanking sequence lengths, i.e. for doublets (XN), triplets (NXN), quadruplets (NXNN), quintuplets (NNXNN) and sextuplets (NNXNNN), where X indicates the position of the substitution in the sequence motif, and N any of the four nucleobases. The results are shown in Table 2, Fig.1, and Supplementary Figs. S1-S4.

**Fig. 1.**
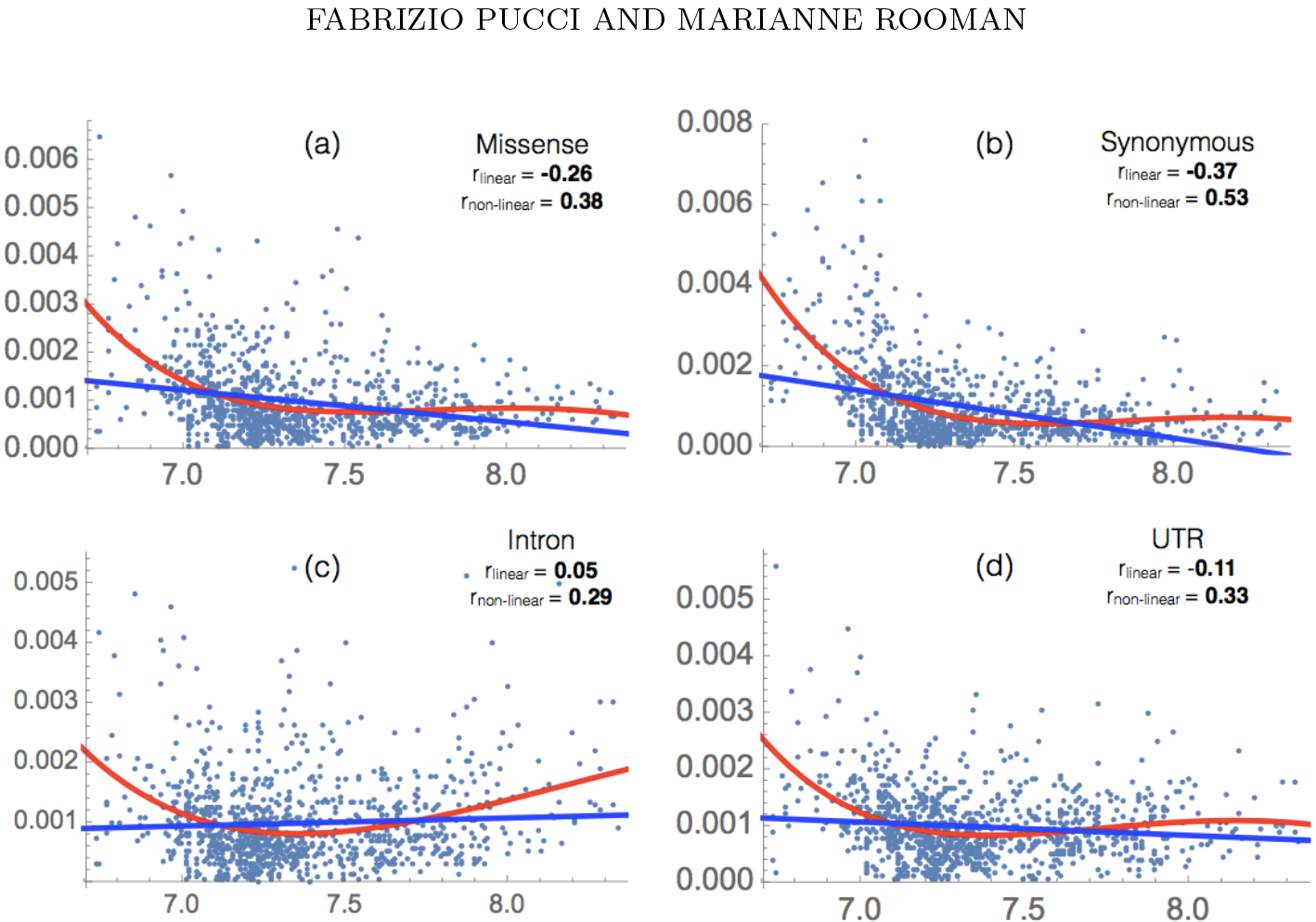
Frequency of observation of SBSs as a function of the vIP (in eV) of nucleobase quintuplets NNXNN, where X indicates the SBS and N any base, for missense, synonymous, intron and UTR mutations. Both the linear (blue) and polynomial (red) regression lines are drawn. The linear (*r*_linear_) and non-linear (*r*_non-linear_) correlation coefficients between the SBSs frequency and the vIP of the quintuplets are indicated.

**Table 2.**
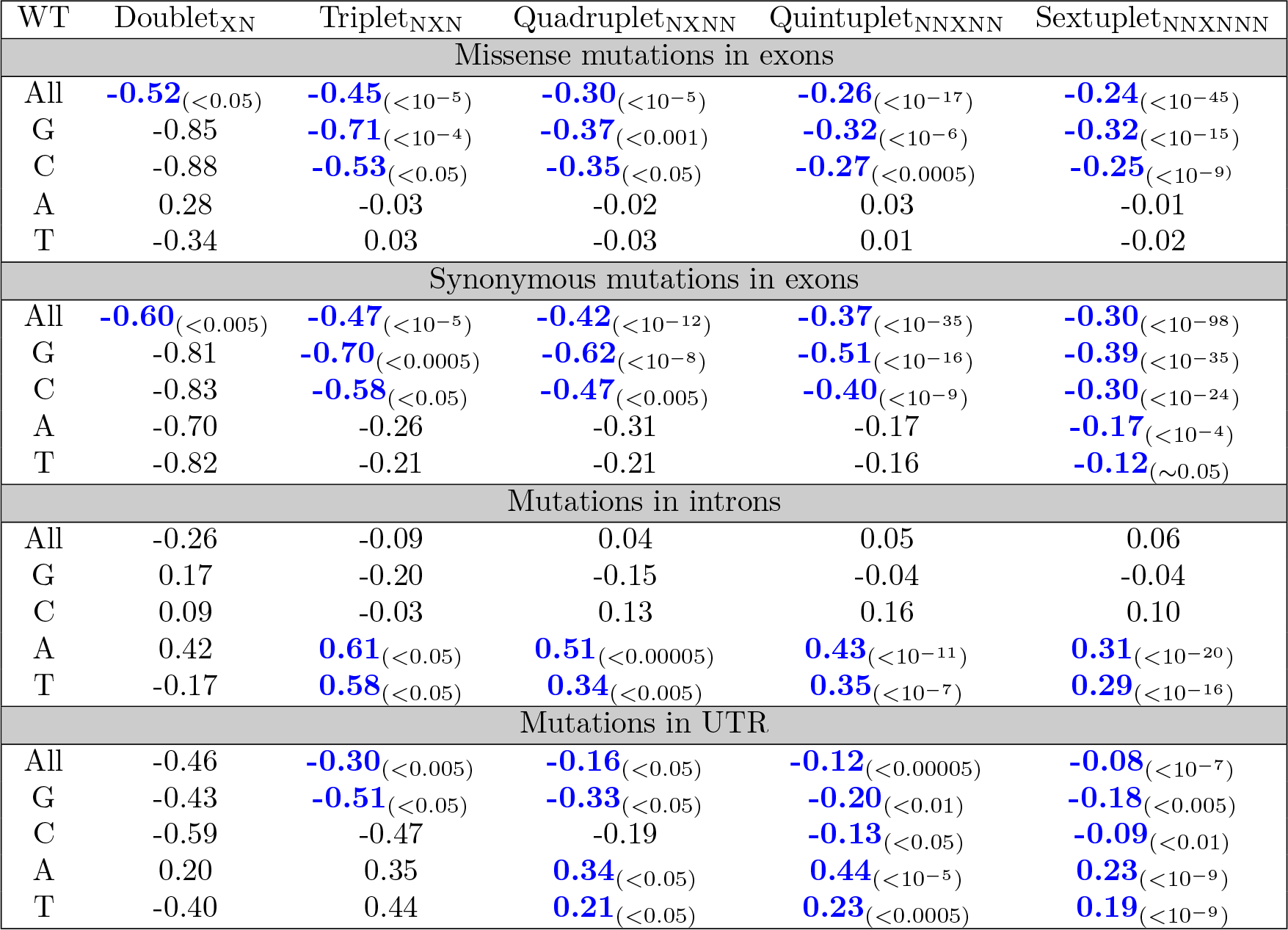
Linear correlation coefficient between the vIPs of the wild-type nucleotides and their flanking base sequences and the associated SBS frequency. The *P*-values are reported in parentheses and the correlation coefficients that are statistical significant are written in blue.

Let us first focus on the behavior displayed by SBSs associated to synonymous and missense mutations occurring in exons. When all possible substitutions are considered together, there is a clear, statistically significant, effect of the frequency modulation of the SBSs by the vIP of the wild-type nucleobase and its neighboring sequence. This “non-local” effect is already visible at the dinucleotide level, and clearly extends to longer sequence motifs. Indeed, for the doublets, the linear correlation coefficient between SBS frequency and vIP values is equal to −0.55 and −0.64 for the missense and synonymous substitutions, respectively. This correlation smoothly decreases if one takes more flanking residues into account, to −0.24 and −0.30 for the sextuplets, while becoming even more statistically significant. In other words, the lower the vIP of the sequence in the vicinity of the SBS, the higher the probability of the middle nucleotide to be substituted. Different biophysical mechanisms are likely to be responsible for this behavior, which will be discussed in the next section.

Interestingly, if we focus on the correlation between vIP and SBS frequency separately for each of the four wild-type nucleotides, we see that the global anticorrelation is mainly due to the sequence motifs with a substituted guanine or cytosine, and not to those with a mutated adenine or thymine. We also observe here a difference between missense and synonymous mutations: the correlation coefficient between the sextuplet vIPs and SBS frequencies of motifs with a substituted adenine or thymine are statistically significant only for the synonymous mutations and not for the missense ones.

The results for the introns differ completely from those obtained for the exons. At first sight, when considering all types of SBSs together, no significant correlation is observed between the vIP of the wild-type sequence motifs and the SBS frequencies. However, when computing the correlation coefficients per substituted nucleotide, the picture becomes different. There is still no correlation for G and C substitutions, but a statistically significant correlation appears for A and T SBSs, for triplets up to sextuplets. Surprisingly, the correlation coefficients are positive and thus, SBSs tend to occur when the wild-type base and its flanking sequence have higher vIP. A more precise analysis, considering the non-linear correlation line depicted in Fig. 1c, shows that SBSs tend to occur when the wild-type base and its flanking sequence have a high vIP, but also when they have a low vIP. This result shows a clear distinction between intronic and exonic SBSs and leads us to conclude that different mutation mechanisms are involved.

In UTR regions, we have a mixture between intronic and exonic tendencies. If we consider SBSs at G or C nucleobases, the mutational rate is higher for low vIPs, like in the exonic case, whereas for A or T nucleobases, the rate is higher for high vIPs, like in the intronic case. This mixed behavior can be unraveled by making the distinction between UTR-3 and UTR-5 regions: mutations in UTR-3 show a similar behavior to those in introns, while mutations in UTR-5 resemble those in exons. (see Table S2 for the comparison of the correlations in UTR-3 and UTR-5).

### 5’-3’ asymmetriy of the flanking sequence in the vIP/SBS frequency correlations

We analyzed whether the nucleobase sequence flanking an SBS in the 5’ direction is more, or less, informative than the sequence in the 3’ direction. For that purpose, we computed the SBS frequency of the nucleotide sequences 5’-XN-3’, 5’-XNN-3’ and 5’-XNNN-3’ and of the ‘’inverse” sequences 5’-NX-3’, 5’-NNX-3’, 5’-NNNX-3’, as well as the average vIP of these sequence motifs. We also computed the correlation coefficients *r*_XN_.._N_ and *r*_N_.._NX_ between the vIPs and SBS frequencies. The results are given in Table 3.

**Table 3.** 5’-3’ directional asymmetry of the SBS flanking sequences. The linear correlation coefficients that are statistical significant are in blue.

|  | Missense | Synonymous | Intron | UTR |
| --- | --- | --- | --- | --- |
| $\langle \text{vIP}_{\text{XN}} \rangle - \langle \text{vIP}_{\text{NX}} \rangle$ (eV) | -0.019 | -0.16 | -0.041 | -0.035 |
| $\langle \text{vIP}_{\text{XNN}} \rangle - \langle \text{vIP}_{\text{NNX}} \rangle$ (eV) | -0.031 | -0.11 | -0.029 | -0.030 |
| $\langle \text{vIP}_{\text{XNNN}} \rangle - \langle \text{vIP}_{\text{NNNX}} \rangle$ (eV) | -0.044 | -0.061 | -0.041 | -0.032 |
| $r_{\text{XN}} - r_{\text{NX}}$ | -0.18 | -0.47 | -0.30 | -0.27 |
| $r_{\text{XNN}} - r_{\text{NNX}}$ | -0.18 | **-0.35**<sub>(<0.05)</sub> | -0.17 | -0.20 |
| $r_{\text{XNNN}} - r_{\text{NNNX}}$ | **-0.21**<sub>(<0.01)</sub> | **-0.16**<sub>(<0.05)</sub> | **-0.18**<sub>(<0.05)</sub> | **-0.18**<sub>(<0.05)</sub> |

We found a clear directional 5’-3’ asymmetry: the vIPs are on the average lower when the flanking base sequence considered is towards the 3’ terminus. Again, the difference is much larger for synonymous mutations than for missense, intronic or UTR SBSs. For doublets, for example, the average vIP of synonymous mutations is 4 to 10 times larger than for the other SBSs. Moreover, the linear correlation coefficients are always more negative when the flanking sequence is towards the 3’ end rather than towards the 5’ end, especially for synonymous mutations, where the difference in correlation coefficient reaches almost 0.5 for doublets. The 5’-3’ asymmetry can be put in relation with the preferential direction of the charge transport along the DNA base stacking, which seems to be from 5’ to 3’ (41).

### Connection between vIP and pathogenicity of the SBSs

Electron hole transport can be expected to play an important role not only in the mutability of specific DNA sequences but also in the pathogenic effect of SBSs, since these substitutions can interfere, for example, with some repair mechanisms through the modification of the affinity for DNA-processing enzymes involved in DNA repair, or with basic cellular processes such as transcription and replication through the modification of targeted protein-DNA interactions.

The first quantity that we computed to gain insights into these matters is the correlation between the vIP of wild-type bases surrounded by their flanking sequence and the difference in frequency between pathogenic and neutral substitutions (Δ*v*). We considered for this purpose both pathogenic and likely pathogenic annotations as pathogenic, and benign and likely benign annotations as benign. As seen in Table 4, there is a small but significant correlation between vIP values and Δ*v*, especially for transversions and missense mutations. In introns and for synonymous mutations, the correlations are not statistically significant, but this is probably due to the small number of such SBSs in our dataset which prevents us to draw robust conclusions. In summary, pathogenic missense mutations are enriched with low vIP sequences, while it is difficult to conclude for other types of SBSs at this stage.

**Table 4.**
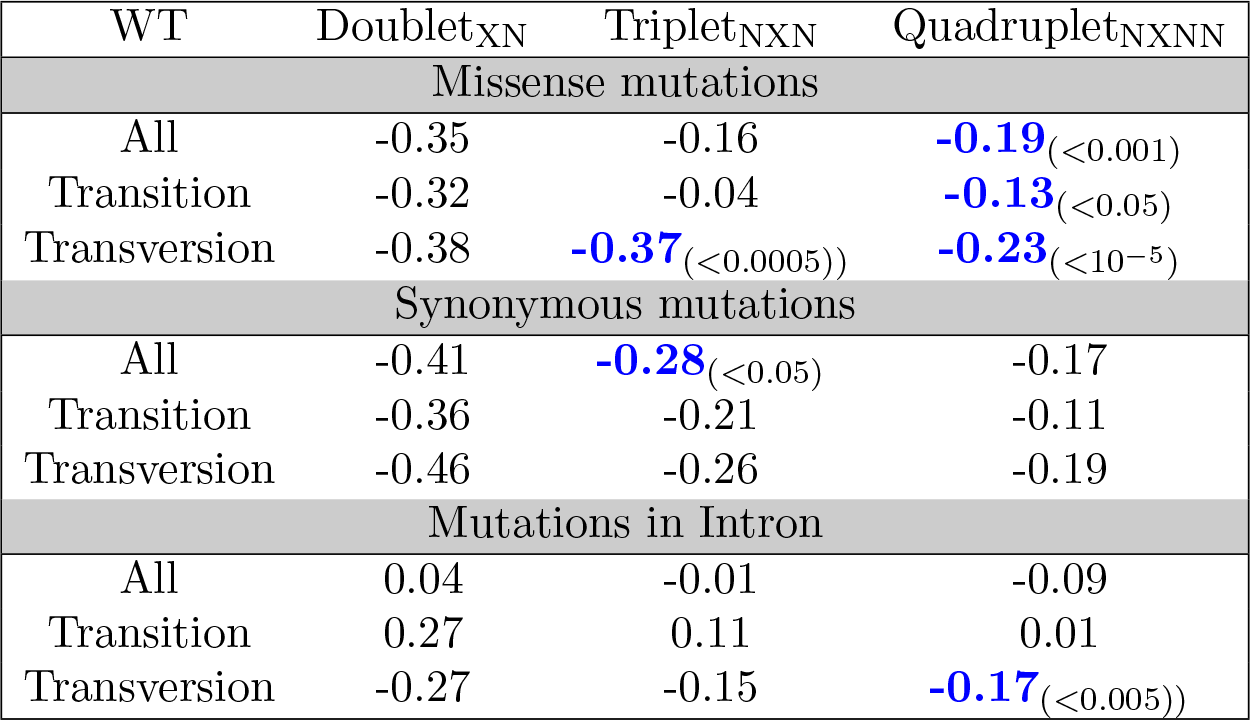
Linear correlation between the vIP of the wild type base surrounded by its flanking sequence and the difference in frequency Δ*v* between pathogenic and benign substitutions.

We also computed the correlation between the absolute values of the change in vIP between the wild type and mutated nucleotides (|ΔvIP|) and Δ*v*. As we can see from Table 5, the correlation is positive and statistically significant and suggests that a larger change in the vIP of the nucleotide base at which the mutation occurs is associated to a higher chance of the mutations to be pathogenic.

**Table 5.**
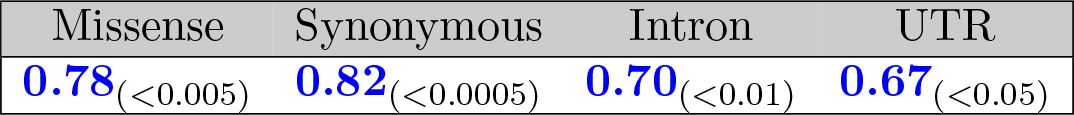
Linear correlation coefficient between the absolute value of the vIP changes upon SBSs (|ΔvIP|) and the difference in frequency Δ*v* between pathogenic and neutral substitutions.

**Table 6.** Experimental vIP values of nucleotide bases (eV) (37; 38).

| Gua | Ade | Cyt | Thy |
| --- | --- | --- | --- |
| 8.20 ($\pm 0.1$) | 8.42 ($\pm 0.1$) | 8.85 ($\pm 0.1$) | 9.10 ($\pm 0.1$) |

Next, we calculated the correlation between the Δ*v* and |ΔvIP| of mutations taking into account the flanking sequences, but no statistically significant correlation was found. This result has several possible explanations. It could be due to the limited accuracy of the vIP calculations and the fact that differences in vIPs have a much larger relative error than vIPs themselves. This is moreover supported by the fact that, when using the calculated vIPs rather than the experimental ones, the correlation drops down to about 0.40. However, this result could also be taken to mean that the observed correlation between Δ*v* and |ΔvIP| is not due to charge migration, but rather to other biophysical phenomena.

## Discussion

Our results show a clear correlation between the vIP of short nucleotide sequences and the frequencies of SBSs in these sequences. However, the type of correlation strongly depends on the gene regions on which we focus our attention, as summarized below.

### In exons, the lower the vIP of a nucleobase sequence segment, the more frequent the SBSs in this segment and vice versa

This effect extends far beyond the dinucleotide units - it is still visible for base sextuplets - and is much stronger for substitutions of G and C bases.

This result indicates the fundamental role played by electron-hole transfer along the DNA stack in mutagenesis processes, which in some cases lead to deleterious phenotypes. The rationale behind this finding is that DNA charge transfer can interfere with different biomolecular mechanisms, triggering SBSs and/or preventing enzymatic-driven DNA repair mechanisms. Indeed, oxidative stress due to physical or chemical agents is likely to cause the extraction of an electron from DNA. Examples of such agents include exposure to ionizing radiation, long-wave ultraviolet (UVA) light, and reactive oxygen species (ROS), just to mention some of them. The hole can then migrate and remain trapped in a minimum of the ionization potentials, and possibly trigger a base substitution. For example, guanine radical cations can react with water to form 8-oxo-guanine, which can trigger G:C→T:A transversions (42), while oxidative damage of the cytosine can undergo deamination and/or dehydration to a poorly repaired uracil, leading after replication to the G:C→A:T transition (for further information on oxidative damage of DNA bases, see (43)).

Quite surprisingly, we found that the correlations are always higher when considering synonymous rather than missense mutations. *A priori*, we did not expect differences in the mechanisms that trigger exon SBSs. It can be argued that this result could be the consequence of codon biases. Alternatively, it could be due to the lethality of some base substitutions that are thus not observed; such mutations are expected to be more often missense than synonymous, as the former modify the proteins and are thus more likely to be deleterious for their function. Or simply, the reason could be that there are more constraints on missense mutations due to their impact on the stability and function of the encoded proteins. Another possible explanation is that the synonymous codon selection is modulated by the constraint of maintaining specific electronic charge properties along the DNA. This would imply the existence of a second code in the DNA, which would lift the codon degenerecy. Other analyses are obviously needed to correctly understand and interpret our results.

Another surprising result is the difference in tendency between exons and introns. For the latter, we found indeed a positive correlation between the vIP of sequence segments and the frequency of substitutions of adenines and thymines in these segments. This means that *in introns, the higher the vIP of a nucleobase sequence, the more frequent the SBSs in the sequence.* This could be viewed as counterintuitive, since it has been suggested that the presence of GGG triplets, which is high near the 5’ termini of introns, could serve as sacrificial anodes to protect the protein-coding part of the genes, and this made us expect a negative correlation (44). However, it is not yet clear what occurs in the deep intronic regions, which mutagenesis mechanisms play a role, and how the correct functioning of the splicing process occurs (45).

An interesting finding is observed in UTRs: while SBSs involving G and C tend to occur when their neighboring sequence has low vIP, the opposite behavior is observed for A and T. However the correlations are much stronger if we consider only 5’-UTRs with an analogous behavior to the synonymous mutations; no correlation is observed in 3’-UTR for G and C substitutions but we found a positive correlation for A and T, a behavior very similar to that of introns (see Table S2).

These UTR regions act to regulate and modulate the protein gene expression at the post-transcriptional level. They contain regulatory sequences that impact the mRNA stability, transport and translational efficiency. Charge transport in these regions can influence a variety of factors and its modification can thus lead to an impact on the normal functioning of the protein synthesis. Note that there are differences between 5’-UTRs and 3’-UTRs in terms of base composition. For example, the former are enriched in G:C and the second in A:T and this could part explain our observations, but this has still to be fully understood.

As clearly visible in Figs 1, the relation between the SBS frequencies and the vIP of the wild-type bases and their flanking sequence does not appear to be linear, and we thus fitted the data using non-linear third-degree polynomials. Of course, there is *a priori* no reason for this correlation to be linear, and this fitting is thus useful to better estimate the probability of mutations of certain DNA sequence stretches, according to their location in the human genome and their vIP values. It would be interesting to scan the entire genome in search for weak regions acting as sinks of electron holes and driving SBSs. Note, however, that the computation of the vIP of longer nucleobase stacks becomes increasingly time-, disk space- and memory-consuming, but cannot be avoided. Indeed, we estimated the vIP of base sequences in two ways, either by calculating directly the vIP of the whole sequence segment by quantum chemistry calculations, or by estimating it as the average of the vIPs of the single nucleobases that compose it. We found that SBS frequencies correlate much better with the vIP values obtained in the first manner. The stacked base structure can indeed not be avoided if we want to explore the behavior of the charge transfer at the quantum level.

Another interesting result is the asymmetry with respect to the flanking sequence when computing the correlations between vIPs and SBS rates. Indeed, the correlation is always better when the mutated base is near the 5’ end of the considered sequence segment rather than near the 3’ terminus. This directional dependence reflects the fact that the physical structure of the DNA stack is asymmetric. For example, the overlap between successive bases differs between 5’-AB-3’ and 5’-BA-3’ configurations (40). This leads in turn to nucleobase stack vIPs which are not symmetric with respect to sequence inversions, and thus to a DNA charge transport with a directional preference. Indeed, even though it remains difficult to test it experimentally, electron hole transfer seems to be more efficient in the 5’-3’ than in the 3’-5’ direction (41). It is thus clear that, if we assume that the charge transport mechanisms are strictly connected to mutagenesis processes, the vIP of the flanking sequence in the 3’ direction after the substituted nucleobase tends to be lower than that of the flanking sequence in the 5’ direction, and to be better correlated with the observed SBS frequency.

Not only mutability appears to be related with electronic properties of DNA, but also pathogenicity. Indeed, we found a weak but statistically significant correlation between the pathogenicity of the SBSs and the vIP values of the wild-type bases and their flanking sequences. It is prevalently observed for missense mutations. Note that this correlation is always larger for transversions than for transitions, and that the former are also more pathogenic in general.

Finally, we found quite a high correlation between the change in vIP between the wild-type and mutant nucleobases in absolute value (|Δ vIP|) and the pathogenicity of the SBSs. However, the correlation disappears when taking also into account the flanking sequences. This can be taken to mean that it is not the vIP that drives the correlation, but other characteristics of the four nucleobases. But it can also be due to the approximations done when calculating vIPs, which are described in the last Methods subsection. Indeed, differences in vIP values are tiny, and inaccuracies in vIPs have therefore a larger effect on ΔvIPs than on the vIPs themselves. More analyses are needed to settle this interesting issue.

Our results led us to conclude that vIPs and the associated DNA charge transfer are not only directly related to the mutation rate, but also to the pathogenicity of the mutations. This effect can be attributed to the modification of biophysical mechanisms such as DNA-protein binding, base excision repair (BER) or nucleotide excision repair (NER) mechanisms (47; 46).

## Conclusion

Since the last two decades, it is becoming increasingly clear that DNA charge transport has a fundamental role in a wide range of biomolecular processes. Sometimes, cells use it for long-range biological sensing and signaling and it is then of vital importance, while in other cases oxidative stress induces electron holes that migrate along the base stack and can, for example, lead to single base mismatches via a wide series of chemical mechanisms, or interfere with the binding of specific proteins.

In this study we focused on the analysis of the SBS rates, and found a clear non-linear correlation with the vIP values of the wild-type bases and their neighboring sequence. Moreover, we observed a different behavior in exons than in introns, in 3’-UTR than in 5’-UTR, for missense mutations than for synonymous mutations, and for G and C bases than for A and T. An 5’-3’ directional asymmetry of the correlation is another indication of the importance of DNA charge transfer in the mutagenesis mechanisms. We also observed a relation between the difference in frequency between pathogenic and neutral mutations and the vIP values, observing that pathogenic mutations are usually embedded in base sequences of lower vIP. A detailed understanding of these results is currently out of reach, but we guess that it would constitute an important step toward the understanding of DNA mutability and how mutations cause pathogenic phenotypes.

Several points remain to be clarified and further explored to get a clearer picture. The first thing that we plan to do is to extend our analysis to longer base sequences, in view of understanding up to which point there is a signal between SBS frequencies and vIP values. Indeed, even though the electron hole trapped in a potential well remains localized in a region composed of 2-4 nucleobases, they can migrate much further away, over distances up to 200 A (16). However, the quantum chemistry calculations of vIP values become more and more challenging and costly when the length of the nucleobase sequence increases, and thus some more approximated methods should probably be considered.

Moreover, vIP values change as a function of the DNA structure (11), so that trapped holes can start migrating again upon (even small) DNA conformational modifications. In this study, we considered only DNA segments in standard B-conformations, but our analysis can easily be extended to A-conformations and other conformations observed in experimental structures. It would also be worthwhile to take into account different nucleotide modifications, such as the 5-methylation of cytosines, as they obviously impact on the vIP values and thus on the derivation of our results.

A last perspective consists in focusing on SBSs occurring in some specific diseases such as cancer and to analyze the link between the observed mutational signatures and vIP values. All the analyses proposed here aim to better understand the mutagenesis processes and how oxidative stress caused by chemical and physical agents affect the the electron-hole transfer in DNA and lead to nucleobase substitutions and pathogenic phenotypes.

## Methods

### Dataset of mutations

The SBS data are extracted from the Single Nucleotide Polymorphism Database (dbSNP) (24). We focused on variants in gene regions, and for all of them, we collected the wild type and mutant nucleotides, the thirty flanking nucleotides, *i.e.* fifteen in the 5’ direction and fifteen in the 3’ direction, the region of the gene in which the mutation is inserted (exon, intron, 5’-UTR, 3’-UTR), the type of substitution (missense, synonymous). If the variant is in an exon region, we moreover identified the exact position of the mutated site with respect to the translated codon.

Our final SBS dataset is then defined as the subset of these SBSs for which the clinical phenotype (benign, likely benign, likely pathogenic, pathogenic, variant of unclear significance,…) is annotated in the ClinVar database (48).

Three quarters of the 190,000 variants from our SBS dataset occur in exon regions, while the remaining are variant in introns (about 25,000) and in UTR regions (30,000 in 3’-UTR and 8,000 in 5’-UTR). Among the exon SBSs, 72% are missense variants and 28% synonymous. Only one third of the variants have an established phenotypic effect (benign, likely benign, likely pathogenic, pathogenic), while the remaining two thirds of the variants have unclear significance (VUS) or other annotations.

### Vertical ionization potentials of nucleobase sequences

For computing the vIP of nucleobase sequences, we used the procedure set up in (11). In a first stage, the four nucleobases Ade, Cyt, Gua, and Thy were considered separately. Their sugar cycle and phosphate group were omitted, and the glycosidic bond was replaced by a hydrogen atom. Their initial geometries were taken fRom The Molecular-Modeling Program Package Insight 2000 (Accelrys Inc.). These Geometries Were Then Optimized At Second-Order MøLler-Plesset Perturbation Theory (Mp2) (29) Using The Split-Valence Basis Set 6-31G* With Added D Polarization Functions On Non-Hydrogen Atoms (30). The Gaussian 09 Program Suite (31) Was Used For These, And All Subsequent, Quantum Chemistry Calculations.

The Energy In Gas Phase Of These Optimized Geometries Was Computed For The Neutral Species And For The Radical Cationic Species, With One Missing Electron. The Calculations For Cationic Molecules Were Performed With Restricted Open-Shell Procedures To Prevent Spin Contamination Problems. The Vip Was Defined As The Difference Between The Energy Of The Cationic And Neutral Species.

In A Second Stage, We Considered All Possible Single-Stranded Nucleobase Stacks In Standard B-Conformation Containing Two, Three Or Four Nucleobases. The Geometries Of The Stacks Were Taken From Early Fiber X-Ray Diffraction Studies (32), Using Insight 2000 (Accelrys Inc.). The Isolated Nucleobases, Individually Optimized At The Mp2/6-31G* Level, Were Superimposed Onto The Original Bases Forming The Stacks, So As To Minimize Their Root Mean Square Deviation Of Atomic Positions Using The U3Best Algorithm (33). The Gas Phase Energies Of These Stacks Of Optimized Nucleobases, For The Neutral And Radical Cationic Species, Were Calculated At Mp2/6-31G* Level, Using The Same Geometry For Both Species. The Vip Was Computed As The Difference Between The Energy Of The Radical Cationic And Neutral Species, Both Adopting The Same Geometry.

Note That We Used Mp2 To Calculate The Vips Rather Than The Hybrid Density Functional Theory Method M06-2X (34) That We Used In (11). Indeed, Although The Vips Of The Single Nucleobases Are Closer To The Experimental Values When Using M06-2X, The Results On Nucleobase Stacks Appear To Be Better When Using Mp2. For Example, The Ggg Triplets Have, As Expected, The Lowest Vip Of All Triplets With Mp2, But Not With M06-2X.

### Approximation In The Vip Computations

We Made Several Approximations When Computing The Vips Of The Nucleobase Stacks. First, We Omitted The Sugar-Phosphate Backbone. Of Course, Their Inclusion Would Require Too Much Computer Time And Memory, But There Are Other Justifications. Calculations On Cyt And Thy Indicated That The Vip Values Change According To The Presence Or Absence Of The Sugar And Phosphate Moieties In Gas Phase, But Much Less In An Aqueous Solvent Due To Screening Effects (35). Moreover, *Ab Initio* Calculations And Experimental Data Showed That The Lowest Ionization Pathway Comes From The Nucleobase Stacks And Not From The Sugar-Phosphate Backbone (36; 35).

The Second Approximation We Made Is To Perform All Calculations In Gas Phase. We Made This Choice To Reduce Computer Time, But Also Because Such Calculations Seem To Suitable For Comparing The Vip Values Of Various Nucle-Obase Stack Sequences. Indeed, Calculations On Individual Nucleobases Highlighted The Effect Of The Solvent In Lowering The Vip Values While Maintaining The Relative Ordering Between The Bases (37; 38). Moreover, Dropping Both The Sugar-Phosphate Backbone And The Solvent Have Opposite Effects That Tend To Cancel Out (35).

We Also Calculated Vips Of Single-Stranded Rather Than Double-Stranded Nucleobase Stacks. This Is Justified By The Fact That Electron Holes Migrate Along A Single Strand And Jump Only Rarely To The Complementary Strand (39). Moreover, We Assumed A Fixed B-Conformation, Although Dna Molecules Have Some Flexibility And The Vips Of Nucleobase Stacks In A- And B-Conformations Have Been Shown To Differ Significantly (11).

### Correlation Coefficients And Statistical Tests

When We Refer To The Linear Correlation Coefficient Of Two Variables X And Y, We Mean The Pearson Correlation Coefficient. We Also Defined A Non-Linear Correlation Coefficient, Obtained Using A Polynomial Function Of Third Degree:

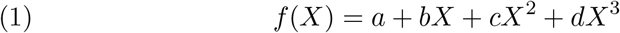

Where The Parameters *A, B, C*, And D Are Identified So As To Minimize The Root Mean Square Deviation Between *F* (*X*) And *Y*. The Non-Linear Correlation Coefficient Is The Pearson Correlation Coefficient Computed Between *F* (*X*) And *Y*.

To Check The Statistical Significance Of Correlations Or Pairs Of Correlations, We Performed A Hypothesis Test On The Bivariate Sample Against The Null Hypothesis. This Was Done By Applying The Fischer Z-Transformation To The Sample, Followed By A T-Test.

## Acknowledgements

We thank Jonathan Velu for help with the SBS dataset construction and Emilie Cauet for interesting discussions. We acknowledge support from the Fund for Scientific Research - FNRS through a PDR research project. FP and MR are FNRS postdoctoral researcher and research director, respectively.

